# Subnetwork-based prognostic biomarkers exhibit performance and robustness superior to gene-based biomarkers in breast cancer

**DOI:** 10.1101/290924

**Authors:** Michal R. Grzadkowski, Syed Haider, Dorota H. Sendorek, Paul C. Boutros

**Author notes:** Corresponding author; 661 University Avenue, Suite 510; Toronto, Ontario, Canada; M5G 0A3. Author Emails: Michal R. Grzadkowski, Syed Haider, Dorota H. Sendorek, Paul C. Boutros.

## Abstract

**Background:** Effective classification of cancer patients into groups with differential survival remains an important and unsolved challenge. Biomarkers have been developed based on mRNA abundance data, but their replicability and clinical utility is modest. Integrating functional information, such as pathway data, has been suggested to improve biomarker performance. To date, however, the advantages of subnetwork-based biomarkers have not been quantified.

**Results:** We deeply sampled the population of prognostic gene-based and subnetwork-based biomarkers in a breast cancer meta-dataset of 4,960 patients. Analysing the performance and robustness of 22,000,000 gene biomarkers and 6,250,000 subnetwork biomarkers across twenty different training:testing cohort partitions of the meta-dataset revealed that subnetwork biomarkers exhibit superior overall performance and higher concordance across partitions. We find evidence of an upper bound for optimal biomarker size of ∼200 genes or ∼100 subnetworks. Additionally, with both biomarker feature types, larger biomarkers tend to show less consistency in performance across partitions, suggestive of over-fitting. Finally, an evaluation of varying training cohort sizes quantifies the effects of training cohort size.

**Conclusions:** Many groups are developing techniques for exploiting network-based representations of biological pathways to characterize cancer and other diseases. By considering the distribution of gene- and subnetwork-based biomarkers, we show that pathway data improves performance and replicability, and that smaller biomarkers are more robust across patient cohorts. These insights may facilitate development of clinically useful biomarkers.

## Background

Breast cancer continues to be the leading cause of neoplastic incidence and mortality in women worldwide [1]□. A major obstacle to the effective diagnosis and treatment of breast tumours is their molecular heterogeneity, which contributes to widely divergent outcomes [2–4]□. Techniques for accurate classification of breast cancer patients by risk profile or therapeutic response are therefore urgently needed as they enable more efficient allocation of clinical resources. A wide variety of models utilizing the mRNA abundance scores of a select set of genes, commonly referred to as “prognostic biomarkers”, have been developed for this purpose in breast cancer [5–11]□ as well as other tumour types and diseases [12–16]□. Although several prognostic biomarkers have proven sufficiently effective to be successfully commercialized [5,6]□, there is no consensus on the best procedure for identifying useful and robust biomarkers. Furthermore, the persistent difficulty in replicating the reported performance of published prognostic biomarkers has led some to question the value of this approach [17–20]□. A proposed method for improving the efficacy and robustness of biomarkers has been to leverage regulatory pathway information. Traditionally, a prognostic biomarker comprises a set of genes whose individual expression levels serve as input for a scoring algorithm that computes the individual risk levels for a patient sample cohort. However, more recently introduced biomarkers consist of *gene subnetworks*, with each subnetwork being a set of genes identified *a priori* as sharing a common functionality via a separate dataset [21–26]□.

Subnetworks are most commonly derived from protein-protein interaction (PPI) networks since protein network data are relatively easy to obtain and contain information on gene-product interactions that it is believed cannot be inferred from mRNA abundance levels alone [27]□. A s*ubnetwork biomarker* thus differs from a *gene biomarker* in that the expression levels of the genes it uses are first aggregated into a set of subnetwork expression scores before being fed into the risk scoring algorithm. We recently introduced a novel scoring system for subnetwork biomarkers constructed using PPI data [28]□. This “SIMMS node-only” model (hereafter referred to simply as *SIMMS*) first calculates the dependence of patient survival on the mRNA abundance of each gene to get a gene risk factor. It then aggregates the gene-level mRNA abundance values within a subnetwork for each patient by taking their sum weighed by the gene risk factors. Several other subnetwork-based schemes have been proposed, and research in this area is active [29–31].

Several studies have cast doubt on the value of using multi-gene models compared to simpler univariate models by citing the performance of subnetwork biomarkers relative to gene biomarkers [32, 33]□. This raises the question of whether time and resources increasingly designated for the development of novel feature selection algorithms as well as the implementation of increasingly composite data for biomarker discovery would be best reserved for research using a more traditional approach of simply identifying patient outcome-related gene sets (*i.e.* gene biomarkers). If building increasingly complex models using advanced machine learning methods and interaction network data yields only marginal improvements in biomarker performance, then identifying informative groups of genes is clearly a more efficient approach to generating clinically relevant prognostic tools.

To resolve this issue, we examined whether using subnetworks as the building blocks of prognostic biomarkers confers any advantage over orthodox gene-based approaches. The SIMMS model was generalized to score gene biomarkers as well as subnetwork biomarkers and then used to evaluate and compare the performance of 22,000,000 gene biomarkers and 6,250,000 subnetwork biomarkers. This experiment was performed with the same set of biomarkers on twenty different permutations of training and testing patient cohorts to test biomarker robustness. Our systematic evaluation of the behaviour of subnetwork biomarkers and gene biomarkers is an important contribution to determining the worth of network-based approaches in biomarker discovery.

## Results and discussion

### Using SIMMS to measure the performance of gene and subnetwork biomarkers

The prognostic performance of subnetwork biomarkers was evaluated on a 4,960 breast cancer patient meta-dataset using the SIMMS model as fully detailed in Methods [28]□. Briefly, a univariate Cox proportional hazards model is used to estimate the association between mRNA abundance levels and survival information of the training cohort for each gene separately. Next, for each patient, the per-gene coefficients from the model are aggregated into subnetwork risk scores. Then, for each subnetwork biomarker, a multivariate Cox model is fit to the patient risk scores of the subnetworks comprising the biomarker. Finally, the subnetwork coefficients of this model are used to generate patient risk scores for an independent testing cohort by combining them with the mRNA abundance values. A univariate Cox model is fit to the testing cohort (dichotomized by the training cohort’s median risk score) and the resulting hazard ratio is used to assess the biomarker’s prognostic capability.

In order to evaluate the prognostic capability of a given gene biomarker in a manner that is directly comparable to subnetwork biomarker performance, we modified the SIMMS model so that it handles individual genes as subnetworks (*i.e.* each gene is treated as a subnetwork of size one). We refer to this approach as *geneSIMMS*.

### Jackknifing reveals patterns in gene and subnetwork biomarker performance

To study the characteristics of the population of gene biomarkers and subnetwork biomarkers, we randomly sampled 22,000,000 gene biomarkers and 6,250,000 subnetwork biomarkers by utilizing *jackknifing*, a resampling method in which samples are drawn from a population without replacement [34]□. For this purpose, we only used genes for which mRNA abundance scores were available in all eighteen datasets comprising our breast cancer meta-dataset (**Table 1**). To ensure that our set of gene biomarkers contained the same underlying mRNA abundance data as our set of subnetwork biomarkers, all gene biomarkers were drawn from a pool of the 1,500 genes which were included in at least one of the 500 subnetworks from which the subnetwork biomarkers were created. Since gene biomarkers were drawn from a set of 1,500 features while subnetwork biomarkers were drawn from a set of 500 features, the population of all possible gene biomarkers is substantially larger than that of all possible subnetwork biomarkers. This necessitated a larger test set of gene biomarkers to fully capture the null distribution of gene biomarker performance. The prognostic capability of each gene and subnetwork biomarker was estimated using SIMMS on twenty different partitions of the breast cancer meta-dataset, each of which was composed of varying training and testing cohort sizes (**Tables 2-3**). For full details on how the meta-dataset was partitioned, see Methods.

**Table 1:** Breast cancer dataset sizes.

| Source | Number of Samples |
| --- | --- |
| Bild | 158 |
| Chin | 129 |
| Desmedt-1 | 198 |
| Desmedt-2 | 107 |
| Hatzis-1 | 310 |
| Hatzis-2 | 198 |
| Ivshina | 249 |
| Metabric-Training | 996 |
| Metabric-Validation | 992 |
| Miller | 236 |
| Pawitan | 159 |
| Sabatier | 252 |
| Schmidt | 200 |
| Sotiriou | 94 |
| Symmans-JBI | 65 |
| Symmans-MDA | 195 |
| Wang | 286 |
| Zhang | 136 |

**Table 2:**
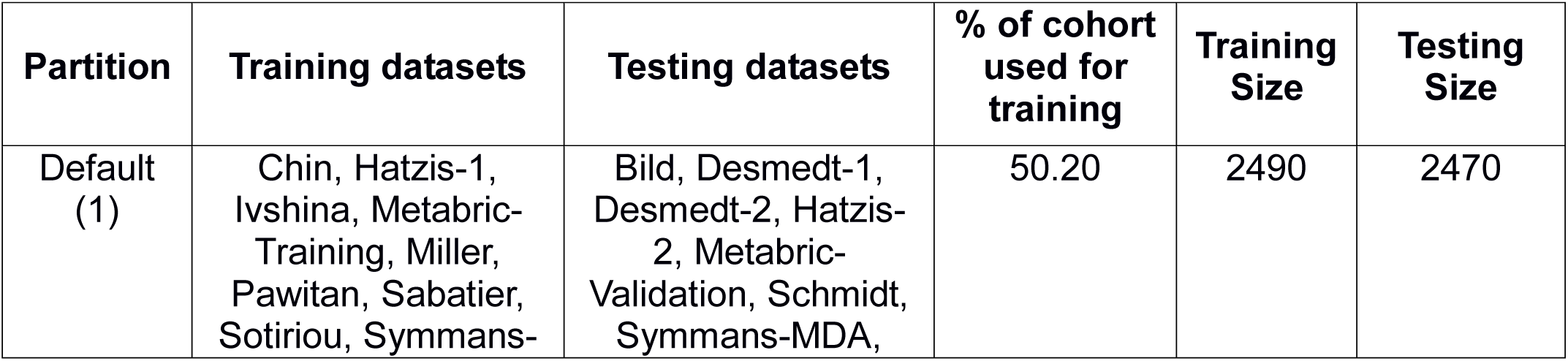
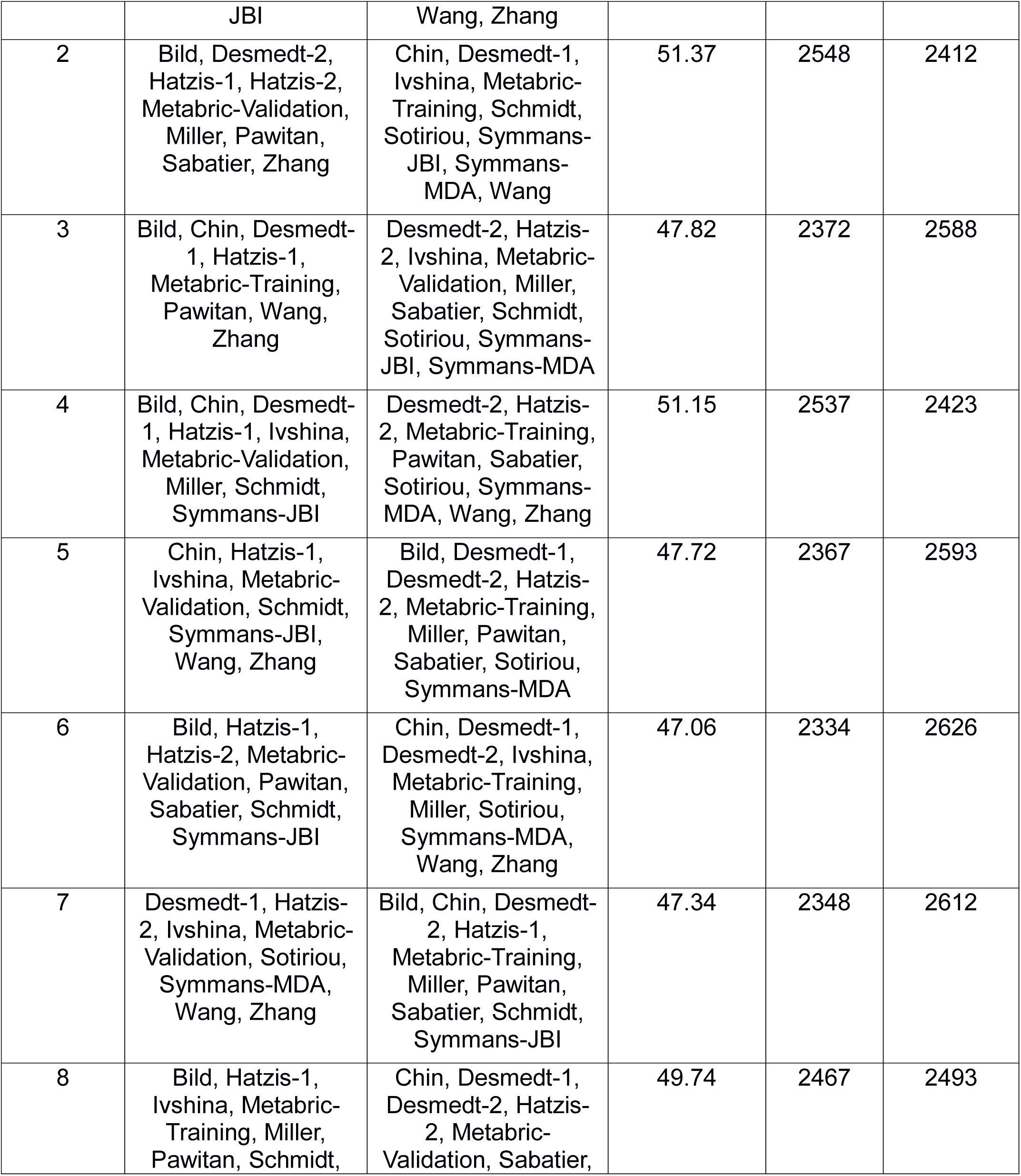
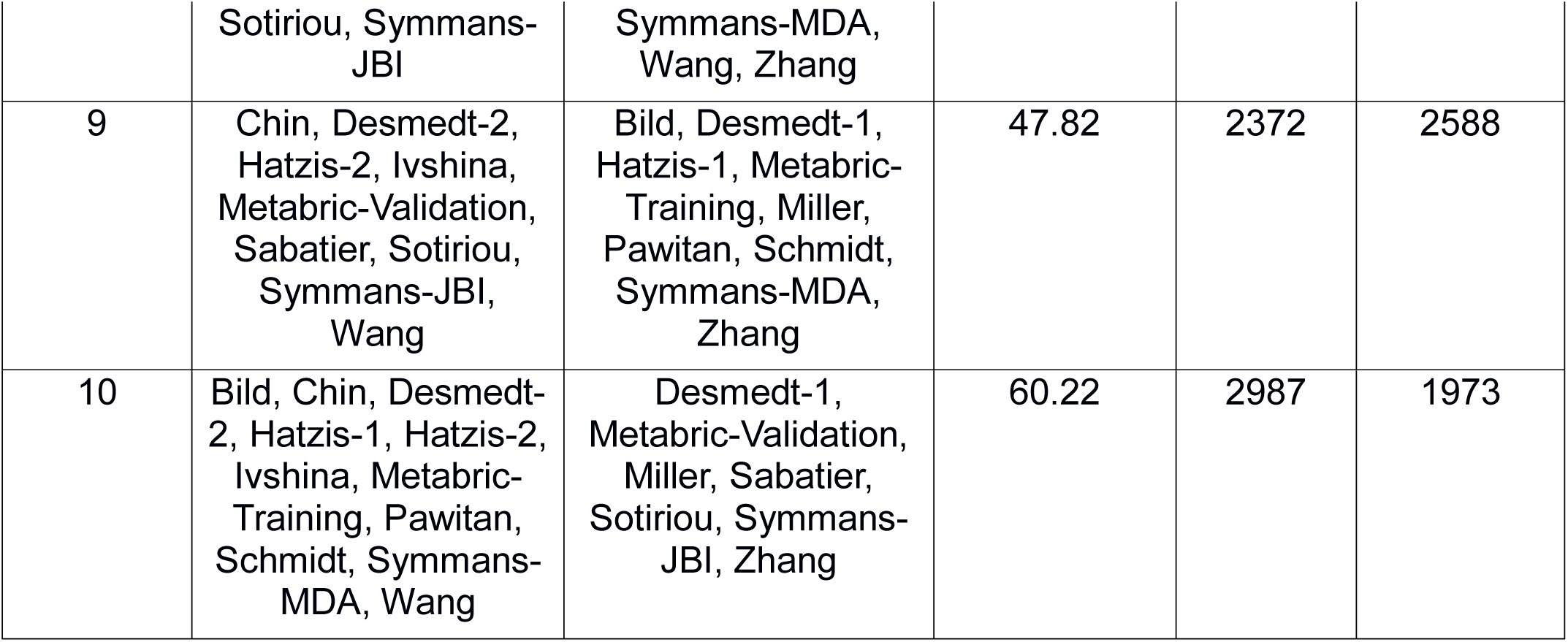
Meta-dataset partitions with larger training cohorts.

Considering only a single meta-dataset partition of equally-sized training and testing cohorts (the ‘default’ partition in **Table 2**), we found that the performance of gene and subnetwork biomarkers is highly sensitive to biomarker size (**Figure 1**). Adding more features to gene biomarkers results in substantial improvements in performance up to ∼100 genes, at which point improvement in performance starts levelling off before peaking at ∼175 genes. After this, larger gene biomarkers show a sharp and consistent decline in prognostic capability, particularly after biomarker size exceeds ∼200 features. On the other hand, the performance of subnetwork biomarkers improves much more gradually with increasing biomarker size, plateauing at around 100 subnetworks.

**Figure 1:**
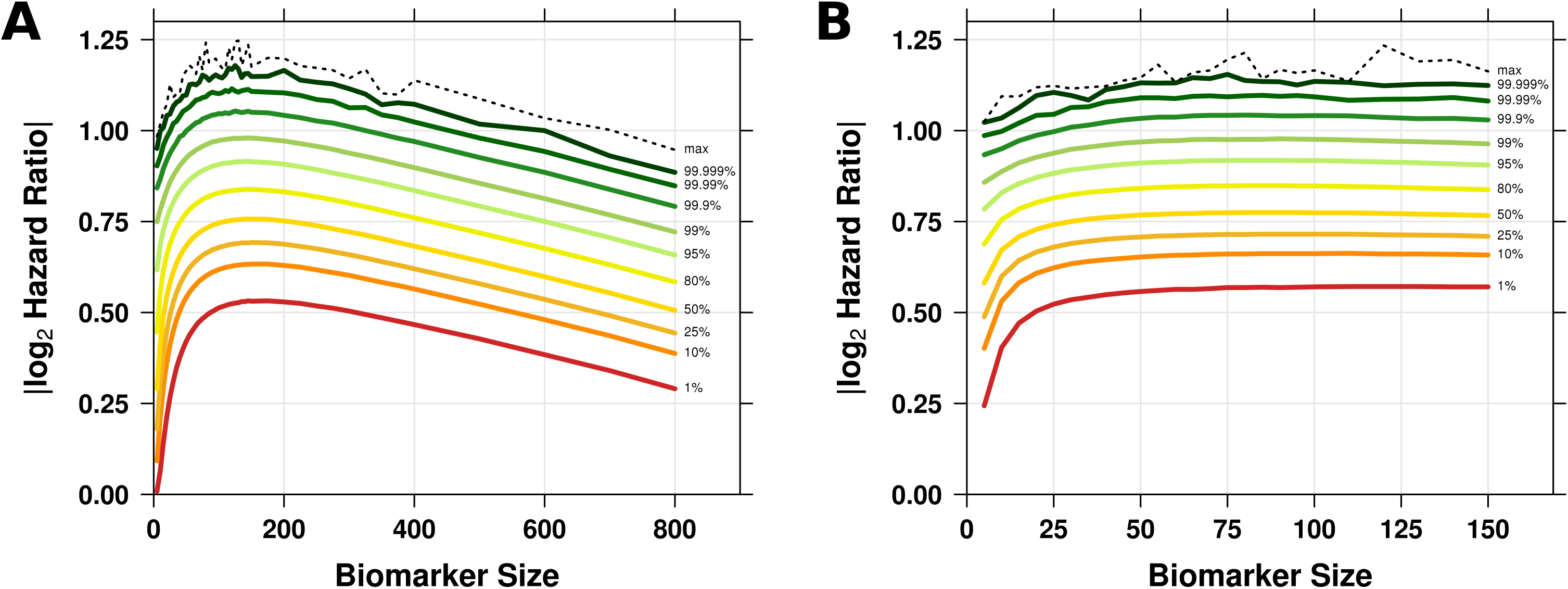
Biomarker performance percentiles on a 50:50 training:testing meta-dataset partition. 250,000 subnetwork biomarkers were sampled at 25 biomarker sizes ranging from 5 to 150 for a total of 6,250,000 subnetwork biomarkers. 500,000 gene biomarkers were sampled using the same pool of genes at 44 biomarker sizes ranging from 5 to 800 for a total of 22,000,000 gene biomarkers. Each biomarker was trained on an initial cohort of 2,490 patients and then applied to dichotomize an independent testing cohort of 2,470 patients using the SIMMS algorithm. The various percentiles of biomarker performance are displayed by biomarker size for **(A)** gene biomarkers and **(B)** subnetwork biomarkers. Performance is defined by the hazard ratio of the Cox model fit to the biomarker dichotomized testing cohort survival data.

### Comparison of gene and subnetwork biomarkers across gene counts

To better contrast the performance of gene and subnetwork biomarkers, we first normalized the sizes of our subnetwork biomarkers to make them more directly comparable to our gene biomarkers. Since a subnetwork biomarker is essentially a set of gene lists, the most straightforward way to perform this normalization is to simply sum the sizes of all these gene lists to get the total number of genes in the biomarker. However, there is gene content overlap in our set of 500 subnetworks so a single subnetwork biomarker may include multiple subnetworks that contain the same gene. Therefore, we additionally considered the number of unique genes in each biomarker. At each gene count, we compared the 95th to 99th percentiles of subnetwork biomarker performance against the 95th to 99th percentiles of gene biomarker performance (**Figure 2**).

**Figure 2:**
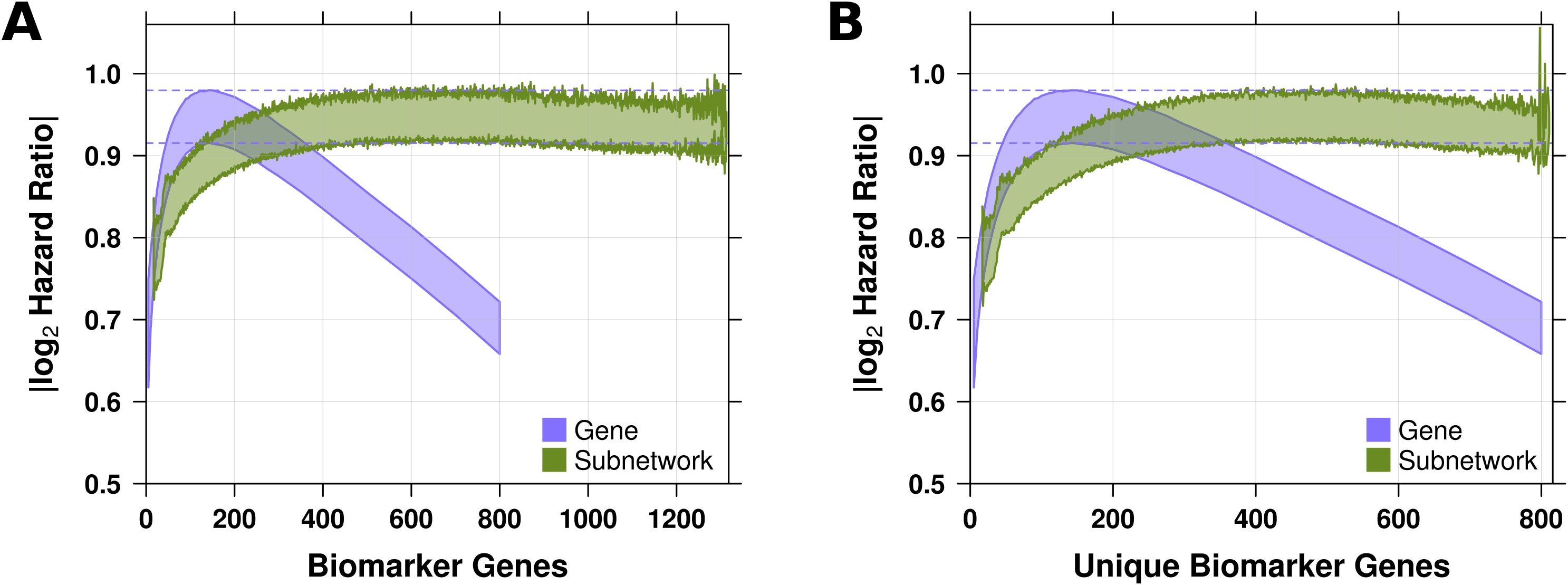
Gene versus subnetwork biomarker performance on 50:50 meta-dataset partition using two biomarker size normalization techniques. For each subnetwork biomarker, we obtained the total number of genes from across all individual subnetworks as well as the number of unique genes. The range between the 95th and 99th percentiles of biomarker performance was re-calculated by grouping biomarkers according to **(A)** the total number of genes and **(B)** the number of unique genes. Note that because the features of a gene biomarker are by definition always a unique set of genes, the total number of genes and the number of unique genes are the same in this case.

The difference in performance between the two gene count normalization techniques is negligible as both gene and subnetwork biomarkers exhibit similar peak performance. Although gene biomarkers require fewer genes to reach optimal prognostic efficiency, they also show a higher propensity for over-fitting at higher gene counts.

### Randomly sampling training:testing meta-dataset partitions reveals effect of training cohort size on biomarker performance and stability

To test the robustness of our finding that biomarker size affects performance, we re-evaluated our biomarker sets on a partition of the meta-dataset with a substantially smaller training cohort at only 36.5% of the total patient number. As seen in **Additional file 1: Supplementary Figure 1**, the general characteristics of biomarker performance with regard to biomarker size remain the same even though the performance of both gene and subnetwork biomarkers decreases slightly overall in this setting. However, subnetwork biomarkers do reach higher peak performance than gene biomarkers, suggesting that they are less sensitive to a suboptimal training environment. To verify these observations, we went on to create eighteen more meta-dataset partitions using random sampling for a total of ten partitions with larger training cohorts and ten partitions with smaller training cohorts.

As first evidenced by our initial finding, both gene and subnetwork biomarkers perform routinely better when trained on the larger patient cohorts (**Figure 3**). In only one dataset partition containing a smaller training cohort do biomarkers consistently outperform those trained on the larger cohorts. For most of the additional eighteen dataset partitions, the performance patterns resemble our initial two partitions. This is particularly true for gene biomarker performance which peaks rapidly at smaller biomarker sizes then drops with increasing biomarker size. Subnetwork biomarker performance peaks slowly, as seen previously, and then either plateaus or undergoes a more gradual decline.

**Figure 3:**
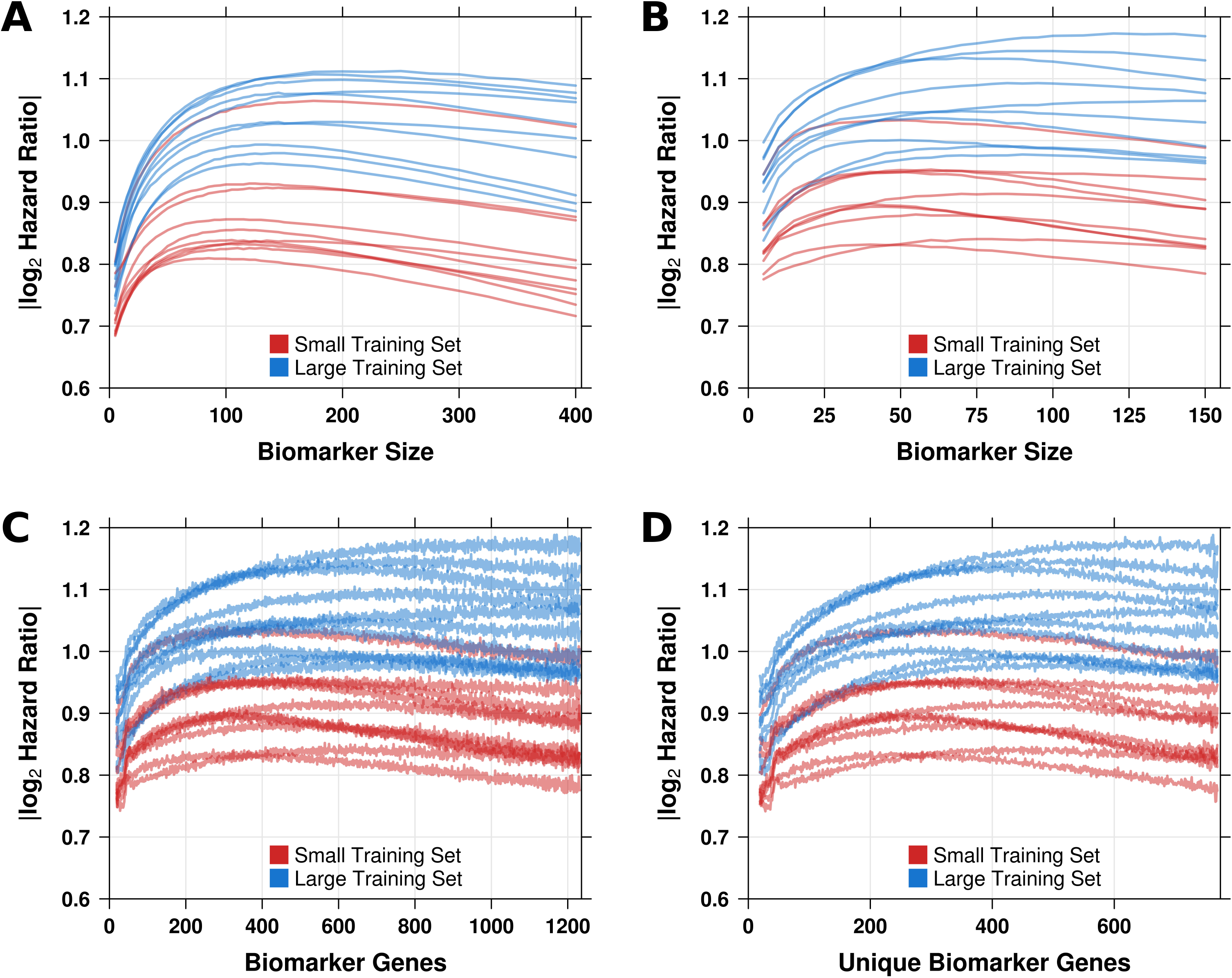
Gene biomarker versus subnetwork biomarker performance across all twenty meta-dataset partitions. Twenty unique partitions of the breast cancer meta-dataset into training and testing cohorts were created such that ten partitions would have at least 47% of the total number of patients to train on (*large training set*) and the remaining ten partitions would have at most 40% of the patients to train on *(small training set*). The performance of 20 million gene biomarkers (the original set of 22 million biomarkers minus those of over 400 genes in size) and the original set of 6,250,000 subnetwork biomarkers was re-evaluated on each of these training:testing cohort partitions. The 99th percentile of **(A)** gene biomarker performance and **(B)** subnetwork biomarker performance on each of the meta-dataset partitions by biomarker size, along with the 99th percentile of subnetwork biomarker performance by **(C)** the total number of genes and by **(D)** the number of unique genes.

To quantify biomarker performance stability across partitions we used Lin’s concordance correlation coefficient (CCC) [35, 36]□. The output of this metric ranges from −1, indicating complete discordance in biomarker performance across all partitions, to 1, indicating identical biomarker performance across all partitions. The stability of biomarker performance at individual biomarker sizes is only slightly greater in partitions with larger training cohorts compared to partitions with smaller training cohorts in gene and subnetwork biomarkers. However, when stability is measured using all biomarkers simultaneously this effect gets amplified (**Additional file 2: Supplementary Figure 2**). More significantly, both classes of biomarkers show the highest performance stability at smaller biomarker sizes, with steep declines in stability as more features are included. This demonstrates that although larger biomarkers may exhibit better prognostic performance, the superiority of individual large biomarkers is much harder to replicate. Therefore, partitioning a dataset to have fewer patients in the training cohort than in the testing cohort has a clear negative effect on both biomarker performance and stability. However, when only small biomarkers are considered, the difference in stability is quite trivial. Since the stability of small gene and subnetwork biomarkers is very high, these biomarkers can be relied upon to have consistent performance when trained on patient cohorts of varying sizes.

### Comparing subnetwork biomarkers to gene biomarkers reveals advantages of subnetworks as features

Finding a link between biomarker performance and feature size prompted us to explore the performance of gene and subnetwork biomarkers in more detail. A clearly defined optimal size was observed for gene biomarkers in all twenty meta-dataset permutations and we used this size as the standard against which subnetwork biomarkers would be judged. Subnetwork biomarker performance was normalized against peak gene biomarker performance in all twenty dataset permutations (**Figure 4**). This reveals that optimal subnetwork biomarker performance tends to be greater than optimal gene biomarker performance, an effect that is particularly pronounced in partitions with small training cohorts. Furthermore, in the majority of partitions where the performance of subnetwork biomarkers exceeds that of optimally-sized gene biomarkers, it does so at a wide range of feature and gene sizes (**Figure 4** and **Additional file 3: Supplementary Figure 3**). In particular, when considering only these partitions, the approximate biomarker sizes at which subnetwork biomarkers start to outperform optimal gene biomarkers is never greater than 50 features, 300 unique genes or 400 total genes.

**Figure 4:**
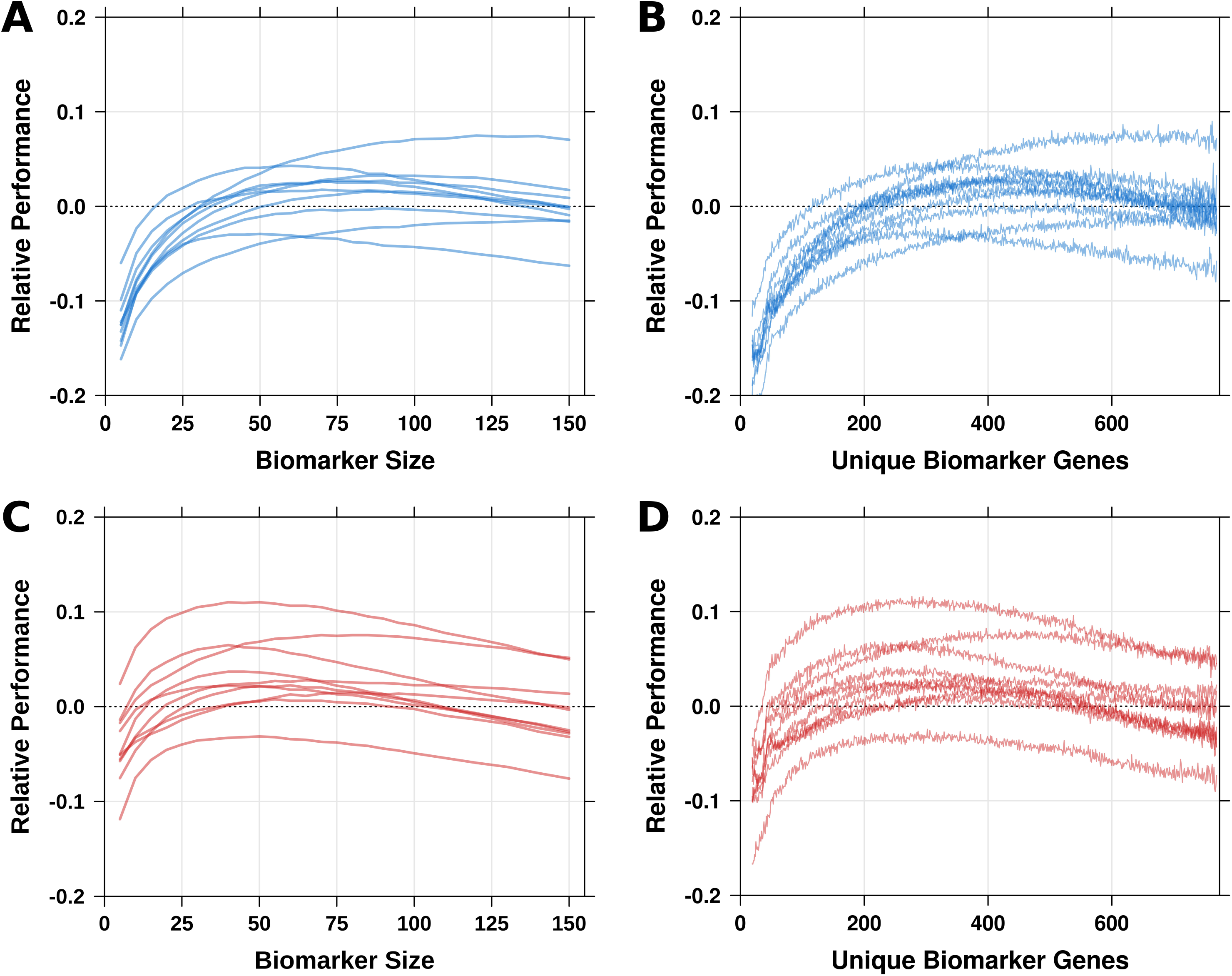
Subnetwork biomarker performance relative to peak gene biomarker performance. On each meta-dataset partition, the peak gene biomarker performance was calculated by taking the maximum of the 99th percentile of jackknifed biomarker performance across all biomarker sizes. The 99th percentile of jackknifed subnetwork biomarker performance at each biomarker size was normalized to this peak on each partition by taking the difference between their Cox model betas. Relative subnetwork biomarker performance on **(A**,**B)** partitions with large training sets (in blue) and on **(C**,**D)** partitions with small training sets (in red) by **(A**,**C)** biomarker size (*i.e.* the number of features) and by **(B**,**D)** the number of unique biomarker genes. For relative subnetwork performance by total number of biomarker genes, see **Additional file 3: Supplementary Figure 3**.

Although the stability of gene biomarker performance is significantly higher than the stability of subnetwork biomarker performance when considering biomarkers of all feature sizes, at comparable gene counts this relationship is inverted (**Figure 5**). The instability of subnetwork biomarkers that contain many large subnetworks obscures the fact that subnetwork biomarkers consisting of a small or moderate number of modules perform relatively consistently across different training:testing cohort partitions. As discussed above, subnetwork biomarkers tend to outperform even optimal gene biomarkers at fairly small sizes. Subnetwork biomarkers without an exorbitant number of features are, therefore, not only more stable than gene biomarkers but also have greater prognostic capabilities.

**Figure 5:**
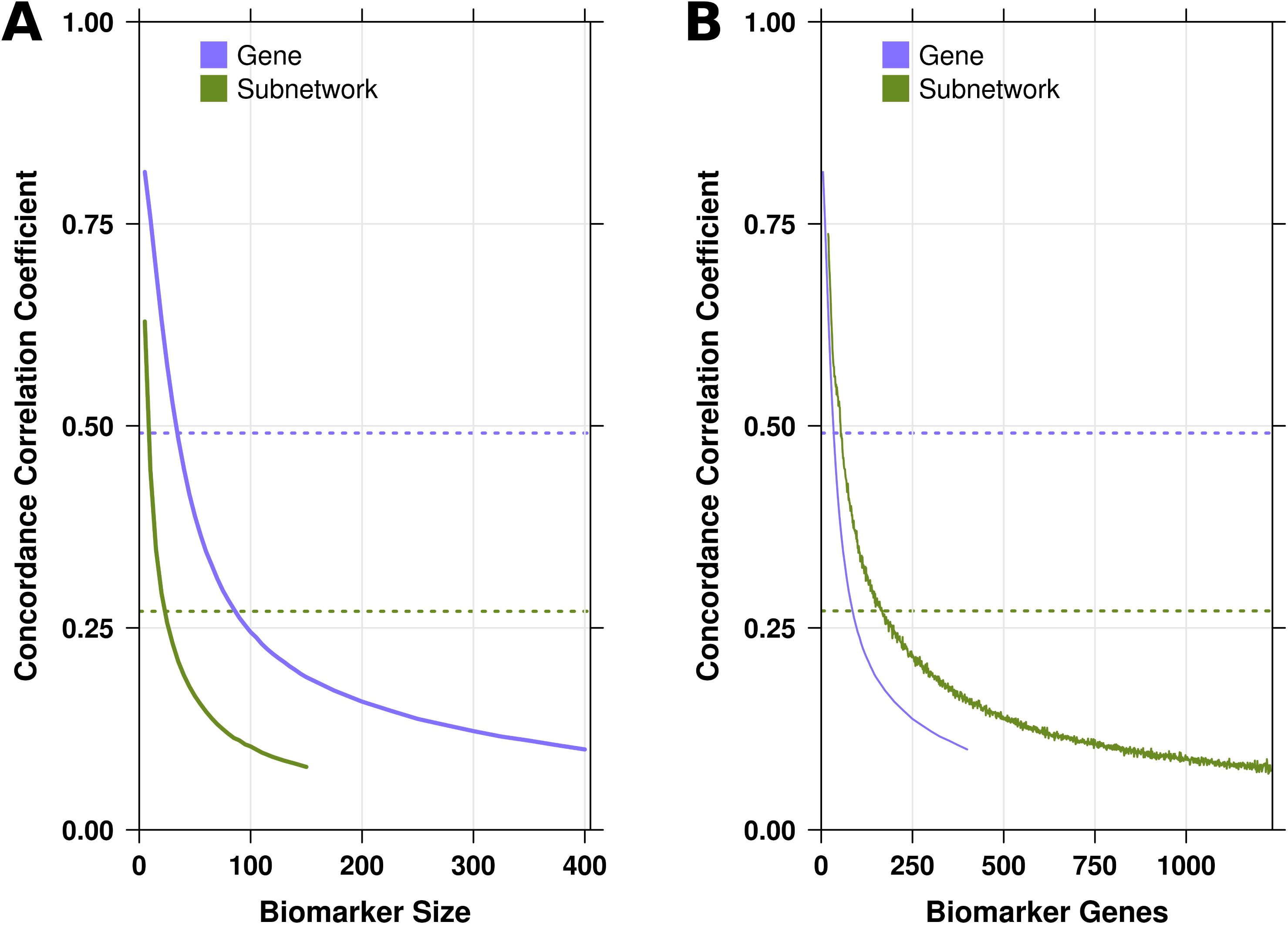
Subnetwork biomarker concordance versus gene biomarker concordance. Lin’s concordance correlation coefficient (CCC) was calculated for gene and subnetwork biomarkers using the twenty meta-dataset partitions as independent observers of biomarker performance. This was done at **(A)** each biomarker size (*i.e.* the number of features) and at **(B)** each total gene count. Dotted lines indicate the CCC calculated for gene and subnetwork biomarkers of all sizes combined.

## Conclusions

In this study, we applied jackknifing to a 4,960 breast cancer patient meta-dataset and identified two key advantages that subnetwork biomarkers have over gene biomarkers. In the majority of our randomly sampled partitions of training and testing patient cohorts, subnetwork biomarkers of various sizes outperform optimally-sized gene biomarkers. Additionally, subnetwork biomarkers show more consistency in their performance across different meta-dataset partitions than gene biomarkers of equal biomarker size (*i.e.* biomarkers containing the same number of features). In other words, a randomly selected subnetwork biomarker can be expected to do a better job of predicting high-risk patients repeatedly across numerous patient cohorts than a randomly selected gene biomarker of comparable size, provided that the number of genes comprising the biomarker does not exceed a relatively high threshold.

The breadth of our analysis also allows us to make several other important observations about prognostic biomarker performance and robustness. Small training cohort sizes have a greater effect on overall biomarker performance than on the consistency of said performance. Furthermore, the performance of large biomarkers is substantially more unstable than that of small biomarkers. Therefore, to optimize biomarker development, it is critical to maximize training cohort size and to minimize biomarker size.

Nevertheless, we recognize there are several caveats to be made regarding our findings that subnetwork biomarkers are superior to gene biomarkers. Because generally gene biomarkers will contain fewer genes than subnetwork biomarkers, and biomarkers with fewer genes are more robust, it is easier to find gene biomarkers with consistently high performance especially since gene biomarker performance peaks at a smaller number of features. Moreover, our study considers only the null distribution of biomarker performance. It is thus possible, albeit highly unlikely, that among the “rare” biomarkers, whose performance falls well above that of those that are randomly sampled, gene biomarkers are preferable to subnetwork biomarkers. Finally, there is a potential limitation in considering only 500 subnetworks and, in particular, only 1500 genes as these may not necessarily be representative of the actual human genome and the approximately 20,000 genes it contains. As information on PPI and gene pathways continues to accumulate, this matter can be thoroughly investigated in the future with access to a more complete and biologically accurate set of data.

Finally, deeply sampling the total biomarker space allows for a comprehensive evaluation of biomarker characteristics that is not attainable through analyses on a smaller number of biomarkers acquired using unsubstantiated feature selection strategies. Since experiment replicability has proven to be a great challenge in the development of clinically useful biomarkers, it is vital to study the performance of biomarkers in a variety of settings, especially using different training and testing cohorts. Jackknifing and meta-dataset permutation are, therefore, two key tools for prognostic biomarker research and can be utilized to great effect in studies hindered by limited sample sizes and data access.

## Methods

### Preparation of mRNA abundance and pathway data

Datasets and subnetworks were prepared as described by Haider *et al* [28]. Raw breast cancer mRNA abundance datasets (**Table 1**) were normalized using the RMA algorithm [37]□ (R package affy v1.28.0) and probes were mapped to Entrez genes using custom CDFs (R packages hgu133ahsentrezgcdf v12.1.0, hgu133bhsentrezgcdf v12.1.0, hgu133plus2hsentrezgcdf v12.1.0, hthgu133ahsentrezgcdf v12.1.0 and hgu95av2hsentrezgcdf v12.1.0). Subnetworks were drawn from the NCI-Nature and BioCarta/Reactome databases and screened such that there was a maximum overlap of 80% between any two subnetworks and each subnetwork consisted of at least three genes. Genes without mRNA abundance values available in all eighteen datasets were removed along with genes that did not appear in any of the subnetworks. This resulted in a pool of 500 subnetworks and 1500 genes.

### SIMMS and geneSIMMS

The original SIMMS node-only model trains a set of subnetworks on a cohort of patients to obtain a “module dysregulation score” (MDS) for each subnetwork. For each gene *g*, the training cohort is median dichotomized by the mRNA abundance value of *g*. A univariate Cox proportional hazards model is then fit to the dichotomization by gene *g* using training cohort survival information and a hazard ratio HR_*g*_ is obtained (R package survival v2.36.14). A risk score for each combination of patient *P* and subnetwork *S* is computed using the formula

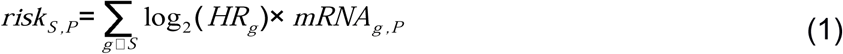

where mRNA_*g,P*_ is the mRNA abundance in patient *P* of gene *g* contained within subnetwork *S*. Next, for each subnetwork biomarker *B*, a multivariate Cox proportional hazards model is fit to its subnetworks’ training cohort’s risk scores, as calculated in **(1)**, and the training cohort survival data to get a Cox beta β_*S*_ for each subnetwork *S* in the biomarker. For each patient *P*, we thus calculate a risk score as measured by *B* using the formula

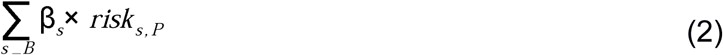

The median risk score is calculated for the training cohort and is then used as the break-point at which to dichotomize the testing cohort. A univariate Cox model is fit to this testing cohort and the risk classification is assessed by the resulting hazard ratio.

The SIMMS model was extended to gene biomarkers by treating each gene as a subnetwork of size one. We refer to this as the geneSIMMS model. The risk score for patient *P* and gene *g* is thus given by

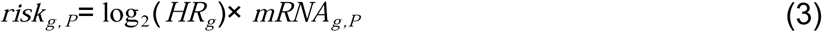

with HR_*g*_ and mRNA_*g,P*_ defined as in **(1)**. For a gene biomarker *B*, a Cox beta β_*g*_ is calculated using a multivariate model for each gene *g* in the biomarker (as is done with the subnetworks in a subnetwork biomarker). For each patient *P*, we thus calculate a risk score as measured by *B* using the formula

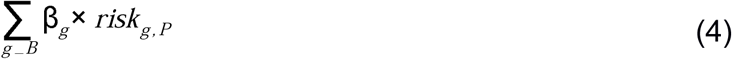

This is followed by patient dichotomization of the testing cohort and then biomarker performance evaluation, same as described above for SIMMS.

### Biomarker jackknifing

Jackknifing is a resampling method in which samples are drawn from a population without replacement [34]□. This technique was applied to estimate the characteristics of the gene and subnetwork biomarker populations. Twenty-five subnetwork biomarker sizes were chosen (all multiples of five between 5 and 100 inclusive and all multiples of ten between 110 and 150 inclusive) along with forty-four gene biomarker sizes (all multiples of five between 5 and 150 inclusive, all multiples of twenty-five between 175 and 400 inclusive and all multiples of one hundred between 500 and 800 inclusive). 250,000 biomarkers were jackknifed at each subnetwork biomarker size for a total of 6,250,000 biomarkers and 500,000 biomarkers were jackknifed at each gene biomarker size for a total of 22,000,000 biomarkers.

The jackknifing was performed to sample more gene biomarkers than subnetwork biomarkers since the pool of potential gene biomarkers is much larger due to there being a greater number of gene features (n = 1500) than subnetwork features (n = 500). Because of the large computational expense of using signatures with a high feature number, gene biomarkers comprising 500 or more genes were only tested on the two initial (default) meta-dataset partitions.

### Normalizing subnetwork biomarker size

A subnetwork biomarker *S* of size *n* consists of a set of modules *M*_*1*_,*M*_*2*_,…*M*_*n*_, each of which is a list of genes. The total number of genes in *S* is therefore

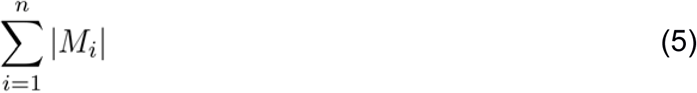

*i.e.* the sum of the sizes of the modules comprising *S*. The number of unique genes in *S* is

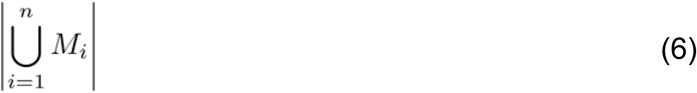

*i.e.* the size of the set of genes appearing in at least one of the modules comprising *S*. A gene biomarker *G* of size *n* consists of a set of *n* unique genes, so the total number of genes and the number of unique genes in *G* both equal *n*.

### Meta-dataset partitioning and concordance correlation calculations

As seen in **Table 1**, our breast cancer meta-dataset consists of eighteen different datasets. To create a partition of this meta-dataset into training and testing patient cohorts, we randomly chose a subset of the eighteen datasets to use for training such that the total number of patients in this subset satisfied some bound and the remaining datasets served as the testing cohort. For partitions with large training cohorts, this bound was at least 47% of the patients in the meta-dataset (**Table 2**). For partitions with small training cohorts this bound was between 33% and 40% of the patients in the meta-dataset (**Table 3**). To ensure random sampling over all partitions satisfying a given bound, we first exhaustively identified all possible partitions and then sampled from that set.

**Table 3:**
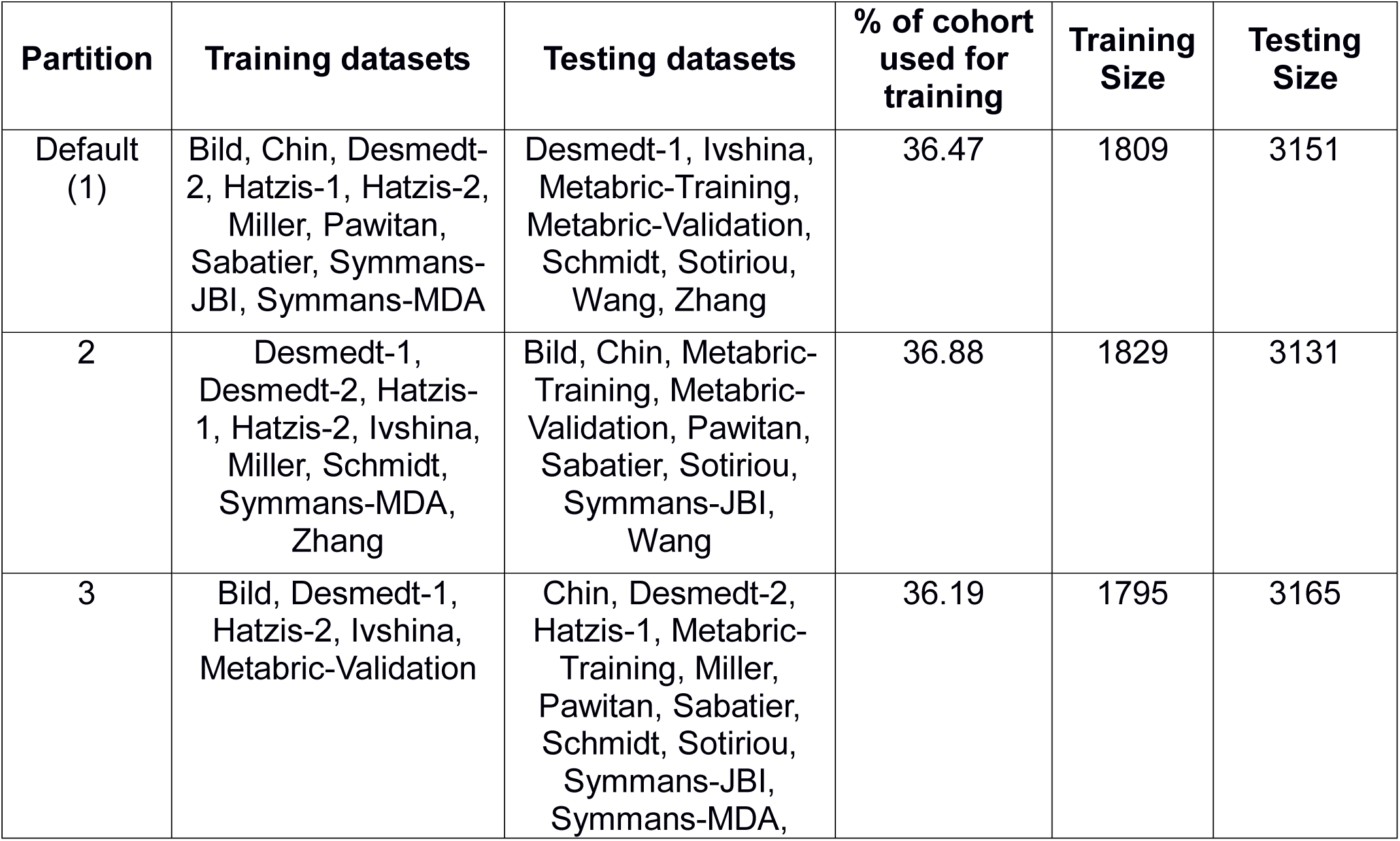
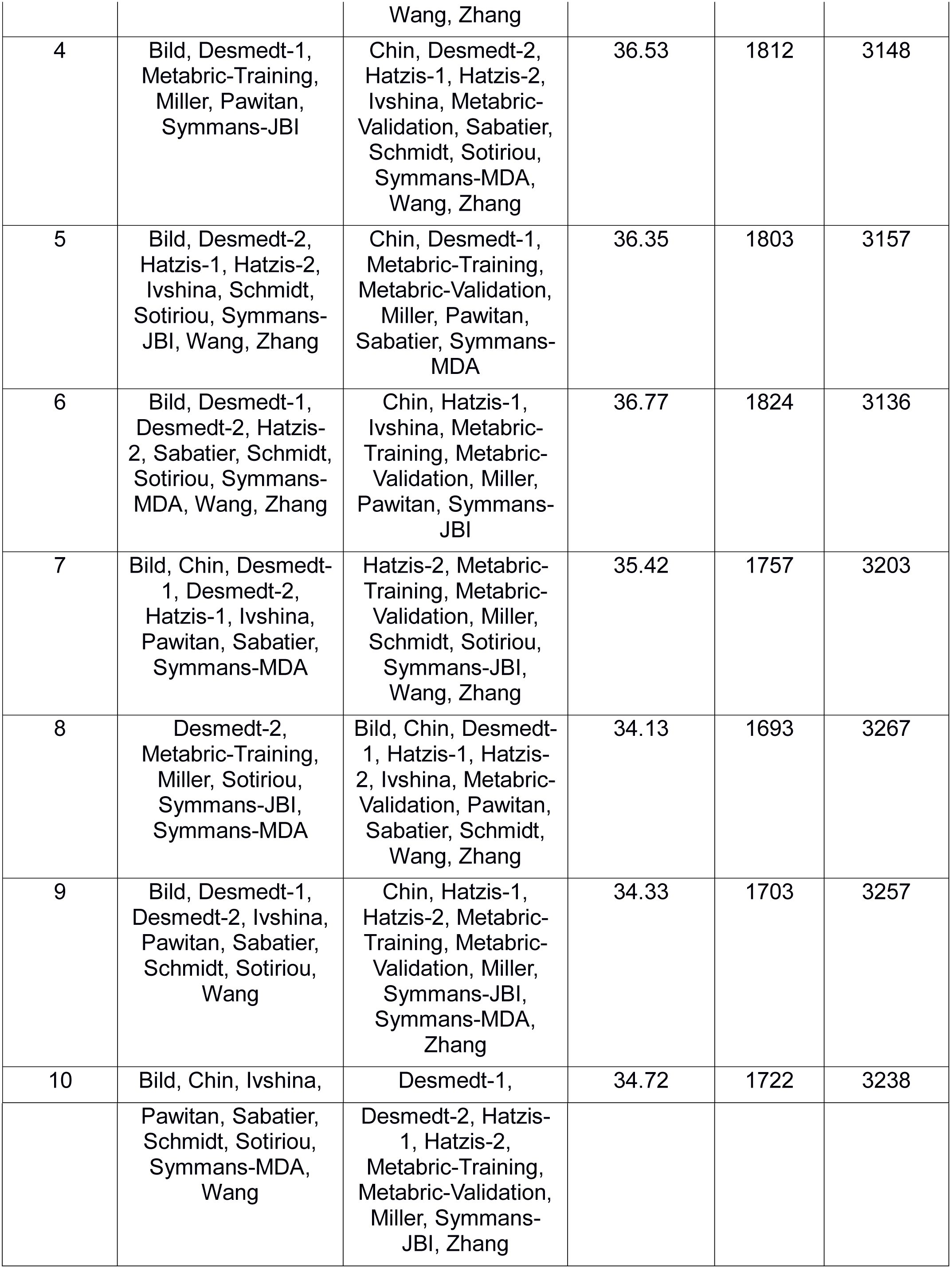
Meta-dataset partitions with smaller training cohorts.

The concordance correlation coefficient for multiple classes was used as first defined and later amended by Lin [35, 36]. Each dataset partition was treated as a class (or observer) in our analysis. Therefore, all gene biomarkers containing 400 or fewer genes and all subnetwork biomarkers had twenty matched observations of their performance as measured by the raw hazard ratio returned by SIMMS or geneSIMMS.

### Comparing subnetwork biomarker performance to peak gene biomarker performance

To determine peak gene biomarker performance on a given meta-dataset partition, we considered the 99th percentile of biomarker performance at each biomarker size as measured by the log-transformed Cox model hazard ratio. The biomarker size which had the highest such percentile of performance defined the optimal size and the corresponding 99th percentile was set as the peak gene biomarker performance for the partition. To compare subnetwork biomarker performance at a given size to this peak, we again took the 99th percentile of performance as measured by the log-transformed hazard ratio and calculated the difference of subnetwork biomarker performance and peak gene biomarker performance to obtain a “relative” performance.

### Visualizations

All figures were created in the R statistical environment (v2.15.3) using the lattice (v0.20-16), latticeExtra (v0.6-24), RColorBrewer (v1.0-5), BPG (v5.6.8) [38] and cluster (v1.14.3) packages.

## List of abbreviations used

CCC: Lin’s concordance correlation coefficient
PPI: protein-protein interaction
SIMMS: subnetwork integration for multi-modal signatures

## Declarations

### Ethics approval and consent to participate

Not applicable

### Consent for publication

Not applicable

### Availability of data and materials

The datasets supporting the conclusions of this article are included within the article and its additional files, as well as obtained from the ArrayExpress, Gene Expression Omnibus (GEO) and European Genome-phenome Archive (EGA) repositories:

- Bild: https://www.ebi.ac.uk/arrayexpress/experiments/E-GEOD-3143
- Chin: https://www.ebi.ac.uk/arrayexpress/experiments/E-TABM-158
- Desmedt-1: https://www.ncbi.nlm.nih.gov/geo/query/acc.cgi?acc=GSE7390
- Desmedt-2: https://www.ncbi.nlm.nih.gov/geo/query/acc.cgi?acc=GSE16446
- Hatzis-1: https://www.ncbi.nlm.nih.gov/geo/query/acc.cgi?acc=GSE25055
- Hatzis-2: https://www.ncbi.nlm.nih.gov/geo/query/acc.cgi?acc=GSE25065
- Ivshina: https://www.ncbi.nlm.nih.gov/geo/query/acc.cgi?acc=GSE4922
- Miller: https://www.ncbi.nlm.nih.gov/geo/query/acc.cgi?acc=GSE3494
- Pawitan: https://www.ncbi.nlm.nih.gov/geo/query/acc.cgi?acc=GSE1456
- Sabatier: https://www.ncbi.nlm.nih.gov/geo/query/acc.cgi?acc=GSE21653
- Schmidt: https://www.ncbi.nlm.nih.gov/geo/query/acc.cgi?acc=GSE11121
- Sotiriou: https://www.ncbi.nlm.nih.gov/geo/query/acc.cgi?acc=GSE6532
- Symmans-JBI: https://www.ncbi.nlm.nih.gov/geo/query/acc.cgi?acc=GSE17700
- Symmans-MDA: https://www.ncbi.nlm.nih.gov/geo/query/acc.cgi?acc=GSE17705
- Wang: https://www.ncbi.nlm.nih.gov/geo/query/acc.cgi?acc=GSE2034
- Zhang: https://www.ncbi.nlm.nih.gov/geo/query/acc.cgi?acc=GSE12093
- Metabric-Training: https://www.ebi.ac.uk/ega/datasets/EGAD00010000210
- Metabric-Validation: https://www.ebi.ac.uk/ega/datasets/EGAD00010000211

## Authors’ contributions

Initiated the project: MRG, PCB Developed the SIMMS model: SH

Extended the SIMMS model to gene biomarkers: MRG

Performed biomarker jackknifing and meta-dataset permutation analysis: MRG

Wrote the first draft of the manuscript: MRG, DHS

Supervised research: PCB

## Acknowledgements

The authors thank all members of the Boutros Lab for their helpful suggestions.

## Additional files

### Additional file 1: Supplementary Figure 1

TIFF 4.5 Mb

**Gene biomarker versus subnetwork biomarker performance using a 37/63 meta-dataset partition**. The performances of the 22,000,000 gene biomarkers and 6,250,000 subnetwork biomarkers sampled were re-evaluated using a smaller training set. From the total 4,960 patient meta-dataset, 1,809 patients were randomly drawn for training and the remaining 3,151 patients were used to test dichotomization efficacy. The various percentiles of biomarker performance using this smaller training cohort by biomarker size for **(A)** gene biomarkers and **(B)** subnetwork biomarkers, along with the range of biomarker performance between the 95th and 99th percentiles by **(C)** the total number of feature genes and by **(D)** the number of unique feature genes for both biomarker types.

### Additional file 2: Supplementary Figure 2

TIFF 4.8 Mb

**Biomarker performance concordance across small training cohorts and large training cohorts**. Lin’s concordance correlation coefficient (CCC) was calculated treating meta-dataset partitions as independent observers of biomarker performance. Partitions were grouped together using the two training cohort size classes (‘small’ and ‘large’) and the CCC for each partition class was calculated for **(A)** gene biomarkers and **(B)** subnetwork biomarkers at each biomarker size and also for subnetwork biomarkers at each **(C)** total gene count and **(D)** unique gene count. Dotted lines indicate the CCC calculated for each partition class using all biomarkers at once.

### Additional file 3: Supplementary Figure 3

TIFF 6.2 Mb

**Subnetwork biomarker performance relative to gene biomarker performance by the total number of genes**. On each meta-dataset partition, the peak gene biomarker performance was calculated by taking the maximum of the 99th percentile of jackknifed biomarker performance across all biomarker sizes. The 99th percentile of jackknifed subnetwork biomarker performance at each biomarker size was normalized to this peak on each partition by taking the difference between the Cox model betas. Relative subnetwork biomarker performance on **(A)** partitions with large training sets and on **(B)** partitions with small training sets by the total number of biomarker genes.

