## Supplementary figures and images for "Subnetwork-based prognostic biomarkers exhibit performance and robustness superior to gene-based biomarkers in breast cancer"

### Supplementary Materials

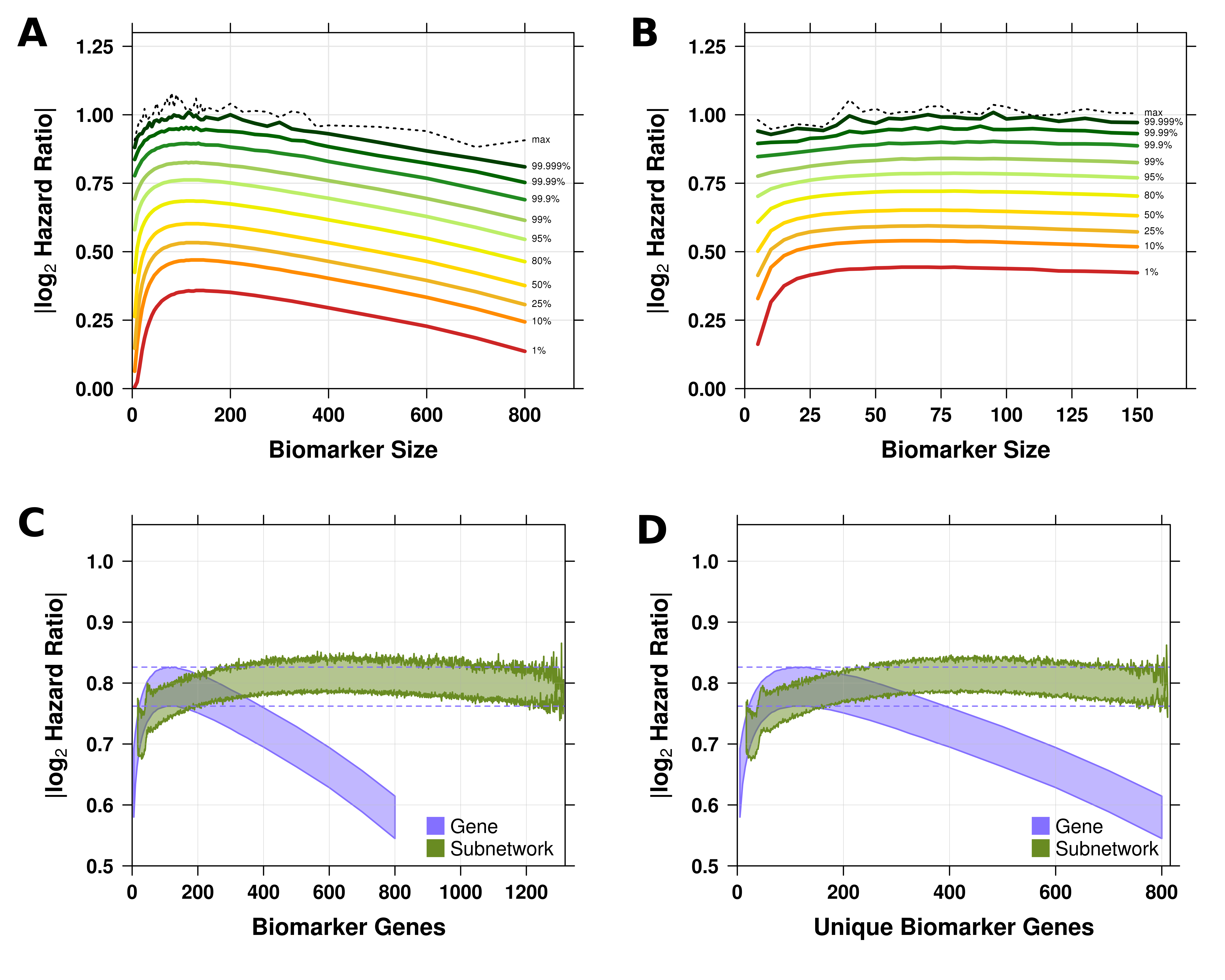

### Supplementary Materials

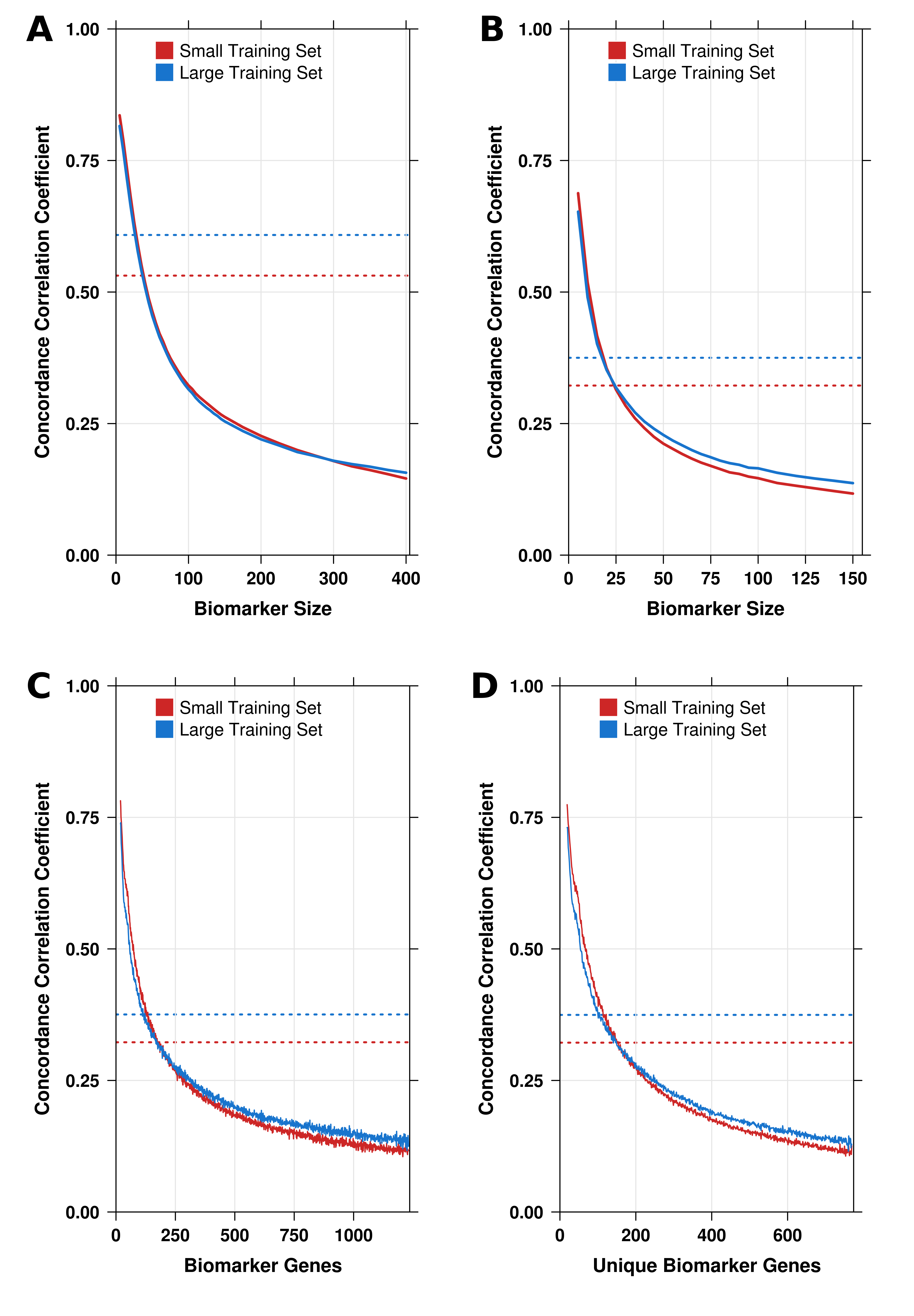

### Supplementary Materials

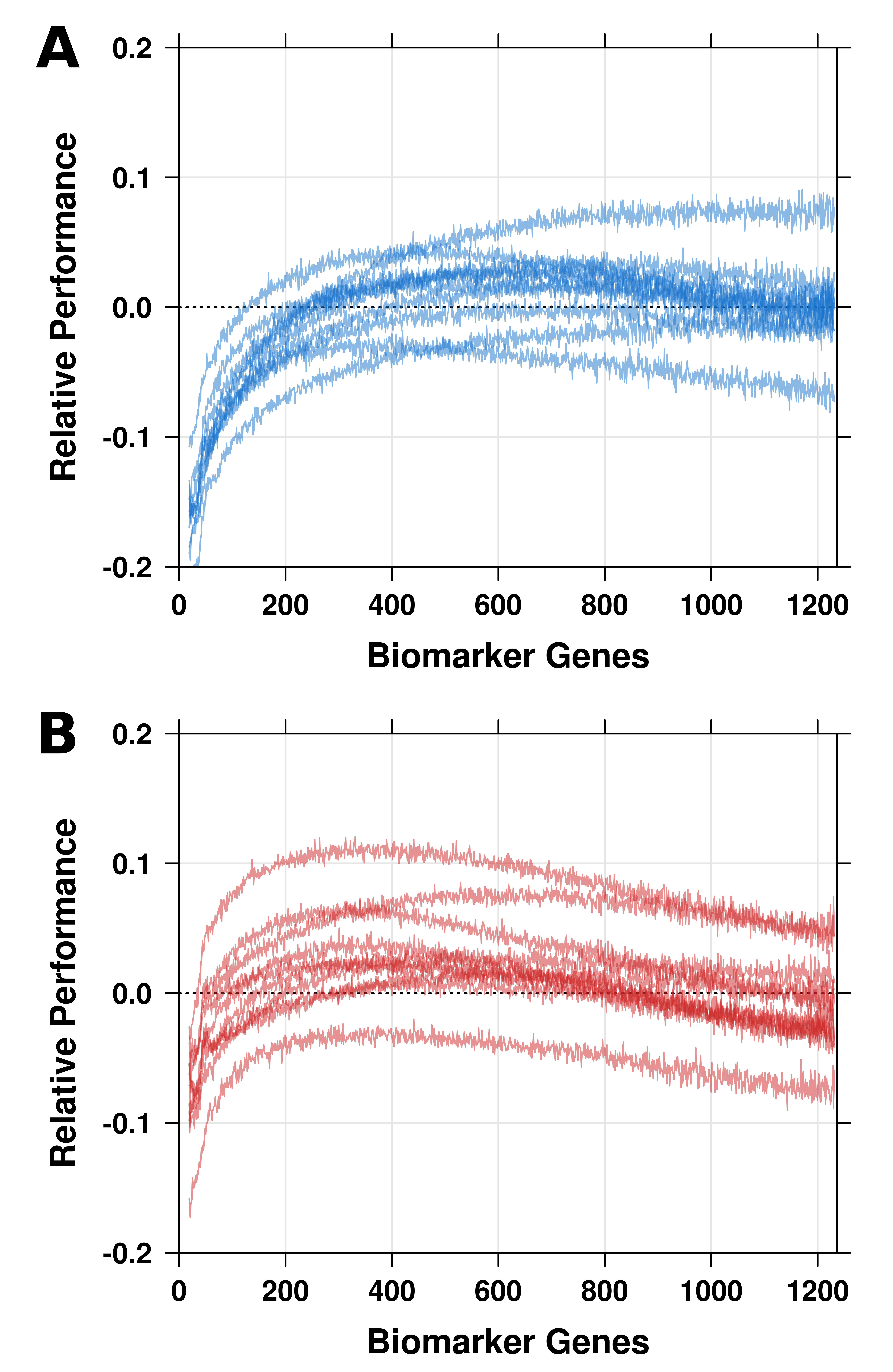
